# Genome-wide CRISPR/Cas9 knockout screen identifies host factors essential for Bovine Parainfluenza Virus Type 3 replication

**DOI:** 10.1101/2025.06.02.657355

**Authors:** Jinyang Hao, Xiaoran Gao, Christine Light, Yuze Sun, Sha Lu, Yu Tian, Xia Gao, Yuan Su, Jie Gao, Xin Huang, Qianyi Zhang, Jinliang Wang, Rong Hai, Wei Hu, Guojun Wang

## Abstract

Bovine Parainfluenza Virus Type 3 (BPIV3) is a leading cause of respiratory illness in cattle and a primary component of Bovine Respiratory Disease Complex (BRDC), resulting in significant economic losses. Understanding the mechanisms of BPIV3 infection, particularly the entry process, is essential for developing effective control measures. Identifying specific host factors that viruses exploit during their life cycle can reveal critical vulnerabilities for potential antiviral targets. We established a genome-wide CRISPR/Cas9 knockout screen in bovine cells to identify host factors involved in viral infections. Our screen identified several key host factors required for BPIV3 infection, including the sialic acid transporter SLC35A1 and the protein LSM12. Further mechanistic analysis revealed that these factors play critical roles at distinct stages of the BPIV3 entry process. These findings not only advance our understanding of how BPIV3 infects host cells, but also identify potential host targets for inhibiting infection and developing novel antiviral strategies.

## 1 Introduction

Bovine Parainfluenza Virus Type 3 (BPIV3) is a leading cause of respiratory illness in cattle worldwide and a primary viral component of the Bovine Respiratory Disease Complex (BRDC), a syndrome that inflicts considerable economic losses on the cattle industry [1, 2]. BPIV3 infection can damage the respiratory tract and compromise the host’s immune system, thereby increasing susceptibility to secondary bacterial infections and exacerbating the severity of disease [1, 3, 4]. The clinical interventions of BPIV3 infection can vary, with young calves and immunocompromised animals often experiencing more severe symptoms [1].

The virus exhibits a global distribution, with distinct genotypes identified in various regions [5]. The widespread prevalence and substantial economic impact of BPIV3 emphasize the urgent need for a comprehensive understanding of its infection mechanisms to facilitate the development of effective control measures.

Like other viruses, BPIV3 relies extensively on the host cell’s machinery to complete its life cycle, including every stage from initial binding to the release of progeny virions [6, 7]. Among those different stages, a thorough understanding of the viral entry mechanisms is particularly crucial, as this initial step dictates the subsequent course of infection and represents a prime target for therapeutic intervention [8, 9]. Identifying the specific host factors that viruses exploit during these processes can reveal critical vulnerabilities, often referred to as their “Achilles’ heels,” which can be targeted for the development of novel antiviral therapies [9–11]. Targeting host factors also offers a promising avenue for broad-spectrum antiviral development and may prove less susceptible to the rapid emergence of viral resistance compared to directly targeting viral proteins [10].

The genome-wide CRISPR/Cas9 knockout screening has been demonstrated to be a powerful tool to identify host factors involved in viral infections due to its high specificity and efficiency in genome editing [8, 10, 12–16]. These loss-of-function screens require the systematic disruption of gene expression across the entire genome to pinpoint genes that are essential for viral replication [8, 12–16]. The CRISPR system offers notable advantages over earlier RNA interference (RNAi) based screens, including higher specificity in gene targeting and a greater likelihood of achieving complete gene knockout [8, 12–16]. The unbiased nature of genome-wide CRISPR screens allows for the discovery of novel and unexpected host factors that might not be implicated through more traditional, targeted approaches.

In this study, we generated an optimized bovine CRISPR knockout library and identified several host factors, such as SLC35A1 and LSM12, required for BPIV3 infection. Our further mechanistic analysis indicated that these factors play critical roles at distinct stages of the BPIV3 entry process. Our results not only advanced our understanding of the BPIV3 entry process, but also identified potential targets for inhibiting infection.

## 2 Results

### 2.1 Development and Construction of the Bovine Genome-wide CRISPR-Cas9 Knockout Library

To perform a genome-wide screen in bovine cells, we first established a stable Cas9-expressing Madin-Darby bovine kidney (MDBK) cell line (M-Cas9) through lentiviral transduction. The efficiency of lentiviral transduction was evaluated by transducing MDBK cells with GFP-expressing lentiviruses at a multiplicity of infection (MOI) of 0.3. Our results showed that the transduction efficiency was 22.5%, indicating that Cas9 expression level in our MDBK cells are suitable for the following screening study (Fig. 1A and Fig. S1A). In the following, we transduced MDBK cells with lentiviruses carrying the Cas9 cassette. We evaluated the integration of the Cas9 gene by polymerase chain reaction (PCR) and Western blotting (Fig. 1A, 1B). Single guide RNA (sgRNA) lentiviruses targeting the randomly selected gene CPT1B were designed to assess the gene-editing capacity of these cell lines. A total of 34 single-cell clones were isolated from screening. Based on the results of the T7E1 and TIDE [17] assays (Fig. 1C, 1D and S1B), we selected an M-Cas9 cell line with the highest efficiency in gene-editing. Subsequently, M-Cas Clone-#21 was chosen to generate a genome-wide CRISPR-Cas9 knockout MDBK library.

**Fig 1.**
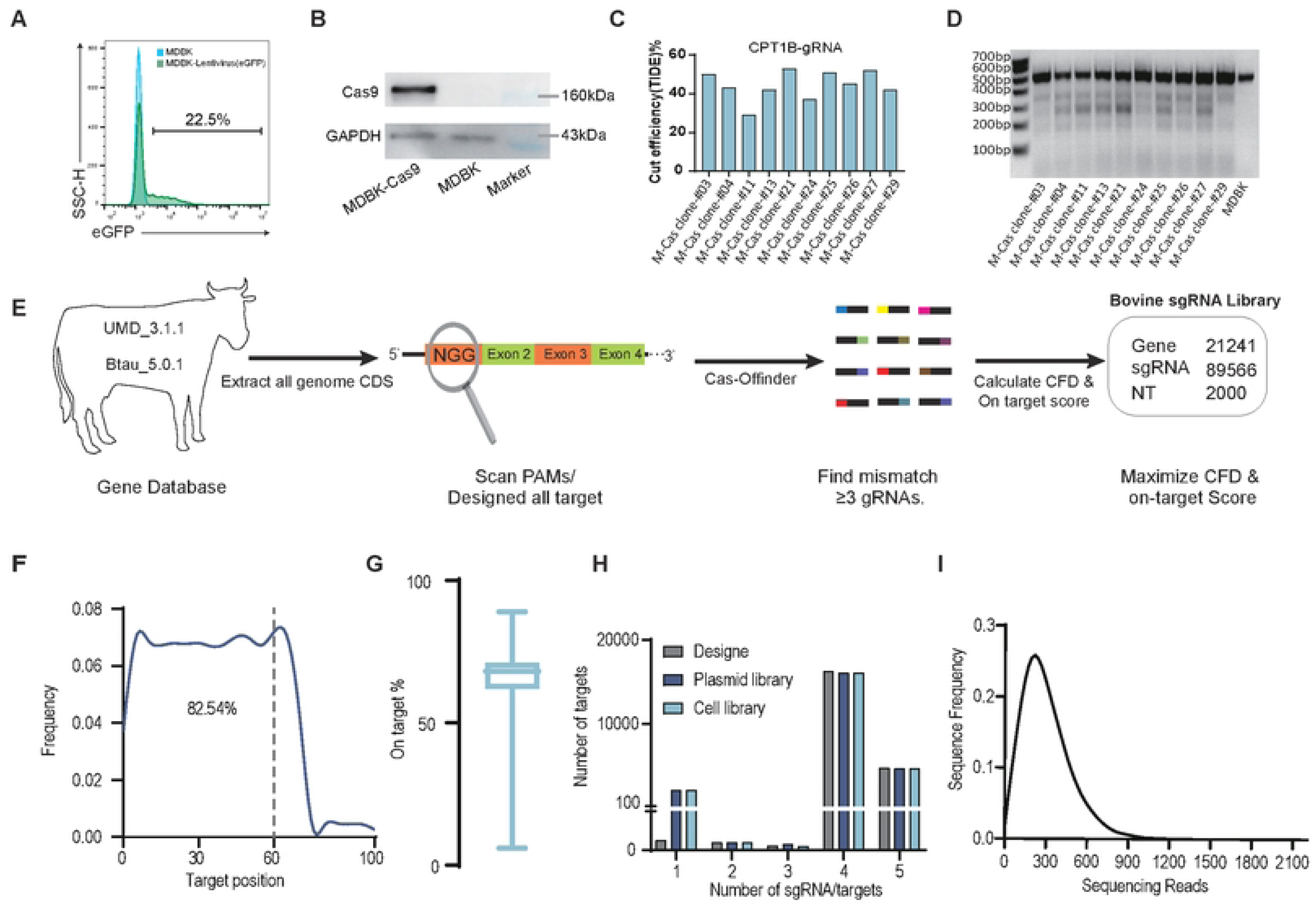
Generation of a bovine gemome-wide lentiviral sgRNA library. (A) MDBK cells transduced with GFP lentivirus and the transduction efficiency measured by FACS. (B) Western blot analysis of Cas9 expression in M-Cas9 cells. (C) Cutting efficiency of M-Cas9-clones in MDBK cells was quantified by TIDE analysis. A sgRNA targeting CPT1B was selected at random to evaluate the cleavage activity of the Cas9 protein expressed in the M-Cas9 cells. (D) whereby ∼200ng of 500bp PCR amplicons of the genomic target from transfected cells were digested by T7 endonuclease I (NEB) and resolved on a 1% agarose gel. (E) This schematic illustrates the pipeline for generating a bovine sgRNA library using CRISPR-Cas9 technology. The process begins with extracting all genome CDS (coding sequences) from the UMD 3.1.1 (University of Maryland r3.1.1) and Btau 5.0.1 (Bos taurus genome assembly version 5.0.1) databases. Subsequently, potential target sites are identified by scanning for PAMs (Protospacer Adjacent Motifs) and designing sgRNAs for all targets. The sgRNA candidates are then subjected to quality control using CasFinder, a tool that calculates the CFD (Cas-OFFinder) and on-target scores to assess off-target effects and on-target efficiency, respectively. The final library comprises 21241 genes, 89566 sgRNAs, and 2000 non-targeting (NT) controls. The workflow ensures the identification of mismatches in ≥3 sgRNAs to maximize the CFD and on-target scores, thereby enhancing the specificity and efficacy of the CRISPR-Cas9 screen. (F) Frequency of the cut positions, calculated as percentages of the full length peptides relative to the 5’ of common coding sequences of targeted genes for all guides. (G) Distribution of cutting efficiency for all guides estimated by crisprScorev.1.1.17. (H) The number of sgRNAs per gene in the genome-wide CRISPR pooled sgRNA library from the designed, plasmid, or mutant cell pools. (I) Cumulative distribution plot of sequencing read frequencies in a CRISPR-Cas9 cell library, showing a peak at approximately 300 reads, indicative of the sequencing depth and coverage across targeted sequences.

To construct a genome-wide knockout library for high-throughput functional genomics research in cattle, using the process for the development of Bovine Genome-wide CRISPR-Cas9 Knockout Library (BovGeCKO) described in Fig. 1E, we integrated the Y chromosome from the btau5.0.1 genome assembly into the UMD3.1.1 genome assembly. For each coding gene target, we used CasFinder [18] to identify all possible 20-nt guide RNAs (gRNAs) adjacent to the canonical protospacer adjacent motif (PAM). gRNAs with three or more mismatches were selected. Each selected sgRNA was then assessed for its on-target and off-target efficiency using the CFD score [19]. For each gene, we chose 4∼5 top candidate guides, which were ranked by cut position and CFD score, to compile a final library consisting of 89,566 guides targeting 21,241 genes, in addition to 2,000 negative controls. In summary, these guides ensured that 82.54% of all targeting guides cut within the first half (0–60% cut position) of common coding sequences (Fig. 1F) and exhibited a median on-target efficiency of 68% (Fig. 1G). Moreover, the plasmid library and BovGeCKO cell collection encompassed all three designed sgRNA sequences for the majority of the targeted loci in the bovine genome (Fig. 1H). The high coverage of the cell library was confirmed by the next-generation sequencing (NGS) (Fig. 1I). Taken together, our results indicate that we have established a highly active and specific BovGeCKO with high application potential for functional genomics research in bovines.

### 2.2 Genome-wide CRISPR-Cas9 screening identified host factors that restrict BPIV3 infection

To enrich a cell population resistant to BPIV3 infection, we infected the BovGeCKO library with BPIV3 virus at a lethal dose, using an MOI of 1 (Fig. 2A). In our BovGeCKO screen, we performed three rounds of BPIV3 infection, using untreated MDBK-Cas9 cells as a negative control to confirm cell death caused by BPIV3 infection in each round. At an MOI of 1, all untreated BPIV3-infected MDBK-Cas9 cells succumbed to infection, whereas a small number of viable cells were detected in the BPIV3-infected BovGeCKO cells. These surviving cells were collected and used for subsequent rounds of BPIV3 infection (Fig. 2A). The sgRNA constructs in the surviving cells were PCR amplified and deep sequenced to identify candidate genes.

**Fig 2.**
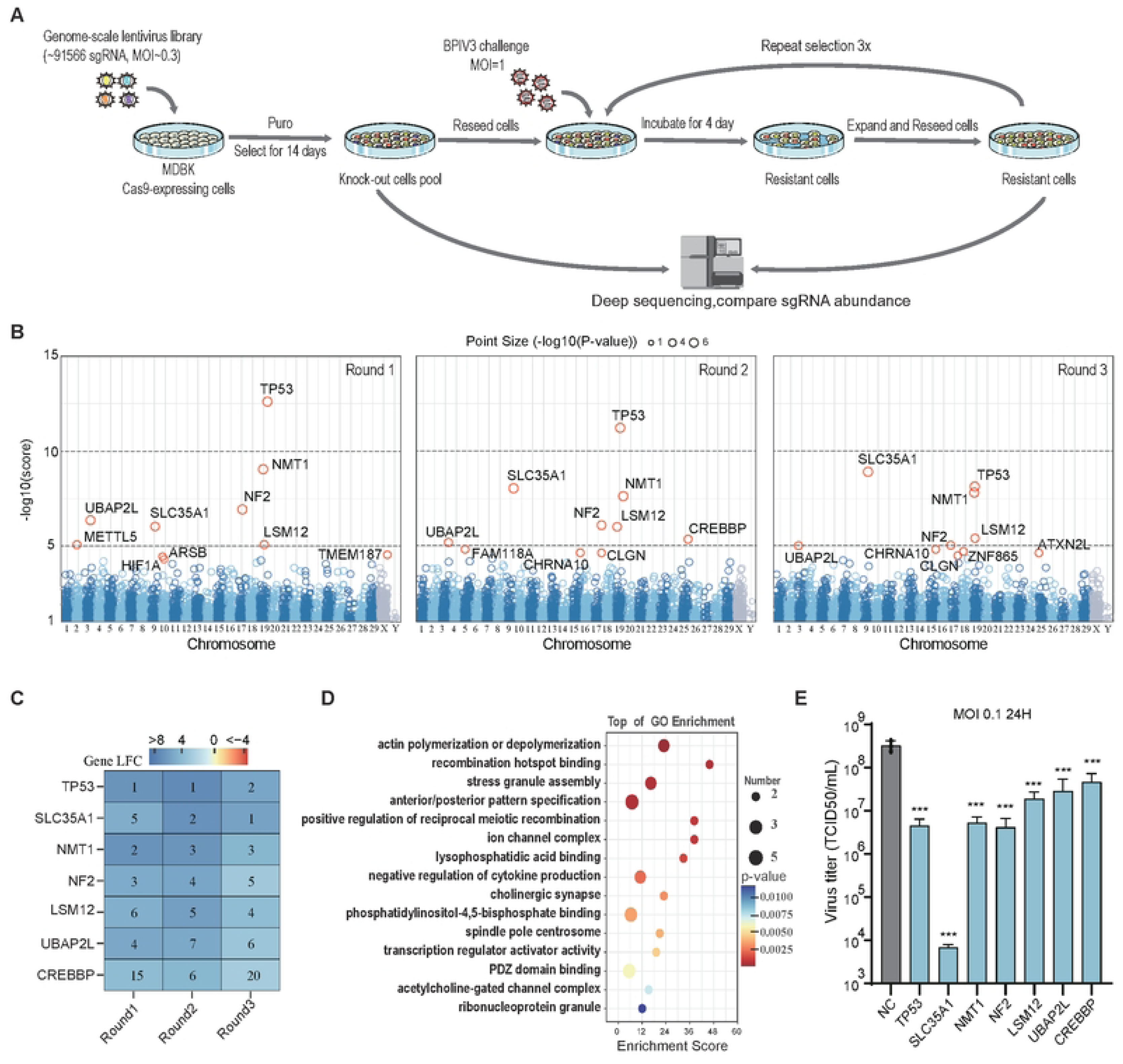
Candidate host genes identified through CRISPR knockout screens. (A) The experimental process involves transducing MDBK cells with a lentiviral library of ∼91,566 sgRNAs targeting the bovine genome at an MOI of 0.3. After 14 days of puromycin selection, cells are challenged with BPIV3 at an MOI of 1. Following a 4-day incubation, resistant cells are expanded, reseeded, and subjected to three rounds of selection to enrich for resistance. Deep sequencing is employed to compare sgRNA abundance, facilitating the identification of key host genes involved in BPIV3 infection. (B) The figure displays scatter plots comparing the robust rank aggregation (RRA) scores of genes positively selected in response to BPIV3 challenges across three rounds. From left to right, the plots represent the first, second, and third rounds of challenges, respectively. These visualizations help assess the consistency of gene rankings and identify key host genes that are consistently important across successive selection rounds. (C) This heatmap illustrates the rankings of selected genes across three rounds of CRISPR-Cas9 screening post-BPIV3 challenge. The numbers within each cell indicate the gene’s rank in the corresponding round, with lower numbers representing higher ranks. The color gradient from blue to red signifies increasing rank, as per the legend. Analyzed using MaGplotR. (D) Gene ontology analysis of 0.5% of the ranked hits from the result of the MaGplotR analysis. (E) BPIV3 replication titers in different gene knockout cells. The cells were infected with BPIV3 at an MOI of 0.01; the supernatants were collected at 24 h post-infection for viral titration in MDBK. The data shown are the means ± SDs of three biological repeats. The two-tailed unpaired t-test was used for the statistical analysis. **, p < 0.002,***, p < 0.001.

Next, we applied the model-based analysis of genome-wide CRISPR-Cas9 knockout (MAGeCK) [20]to identify enriched sgRNAs in surviving cells. A total of 154 genes were identified (p < 0.01, sgRNA hits ≥ 3). After the third round of BPIV3 infection, the most enriched candidate genes, in descending order, were SLC35A1, TP53, NMT1, LSM12, UBAP2L, NF2 and CHRAN10 (Fig. 2B). To further elucidate cellular functions critical for BPIV3 infection in MDBK-Cas9 cells, we compared the sgRNA and gene distributions across the three screening rounds using MaGplotR [21] analysis. This analysis identified the top seven (TOP7) genes that consistently exhibited a fold change greater than 4 logarithms across all three challenge rounds and could be represented by more than 3 independent sgRNAs (Fig. 2C, S2A and Table S2).

Our findings indicated the efficiency of CRISPR-based positive selection screening by reliably identifying potential host factors required for infection. To elucidate the potential biological roles of these candidate genes required for BPIV3 infection, we conducted Gene Ontology (GO) pathway enrichment analyses on the top 0.5% ranked sgRNA targets identified through MaGplotR analysis (Fig. 2D). The analyses revealed significant enrichment of genes involved in actin polymerization and depolymerization, as well as ion channel complex formation, processes that are crucial for various viral activities, including endocytosis, phagocytosis, and organelle dynamics (Table S3). Additionally, we observed enrichment of genes associated with stress granule assembly, negative regulation of cytokine production, and transcription regulator activator activity, along with genes implicated in intracellular signaling pathways and the regulation of cell-intrinsic immunity (Table S3). These results advanced our understanding of the molecular mechanisms required for BPIV3, which were previously unknown.

We next examined the physiological roles of TOP7 genes by evaluating viral replication in cells deficient for each individual TOP7 factor. MDBK monoclonal cell lines with knockouts of the TOP7 genes were generated by puromycin selection and limiting dilution assay, with each knockout confirmed by sequencing the sgRNA target site (Fig. S2B). These results were then compared to viral replication in control cells treated with non-targeting guides. In all cases, depletion of the TOP7 genes led to a substantial reduction in virus titer compared to the control (Fig. 2E). Among these, SLC35A1 and LSM12 exhibited stable and concordant gRNA enrichment, with most sgRNAs showing consistent positive selection across all rounds. (Fig. S2A). Therefore, considering the stability of gRNA enrichment across the three screens, we selected SLC35A1 and LSM12 to further characterize their mechanisms.

### 2.3 SLC35A1 deficiency affects BPIV3 attachment

It is known that the SLC35A1 gene encodes the CMP-sialic acid transporter (CST) necessary for the sialylation of proteins and lipids [15, 22, 23]. Sialic acids are a family of sugars with a nine-carbon backbone that are typically found at the terminal positions of cell-surfaces and secreted molecules [24]. BPIV3 infection of a host cell is initiated by the binding of viral haemagglutinin-neuraminidase(HN) to sialic acid moieties, which are terminal sugars on glycans [25]. Thus, we hypothesize that the SLC35A1 gene plays an important role in helping the binding of BPIV3. We first assessed the levels of virus binding on the cell surface. SLC35A1 knockout (KO) cells exhibited a significant restriction of BPIV3 binding, with only 10% of the cells testing positive for viral infection (Fig. 3A). As a control, we tested the cell surface sialic acid levels in our SLC35A1 KO cells. We used specific lectins to detect 2-3 (Maackia amurensis lectin [MAL]) or 2-6 (Sambucus nigra lectin [SNA]) linked sialic acids on the cell surface. Our confocal microscopy analyses revealed that SLC35A1 KOs lacked binding for both types of lectins compared to vector control cells, indicating a loss of cell-surface sialic acid in the absence of SLC35A1 (Fig. 3B and 3C). To investigate the effect of sialylation of glycans on viral infection, we inhibited the synthesis of sialoglycans using 3Fax-Neu5Ac, a fluorinated sialic acid analogue, in MDBK cells. We found that 3Fax-Neu5Ac effectively depleted α2,3/α2,6-linked sialic acids in MDBK cells (Fig. 3D), and treatment of MDBK cells with 3Fax-Neu5Ac resulted in a 74-fold suppression of BPIV3 replication compared to DMSO-treated cells (Fig. 3E, F).

**Fig 3.**
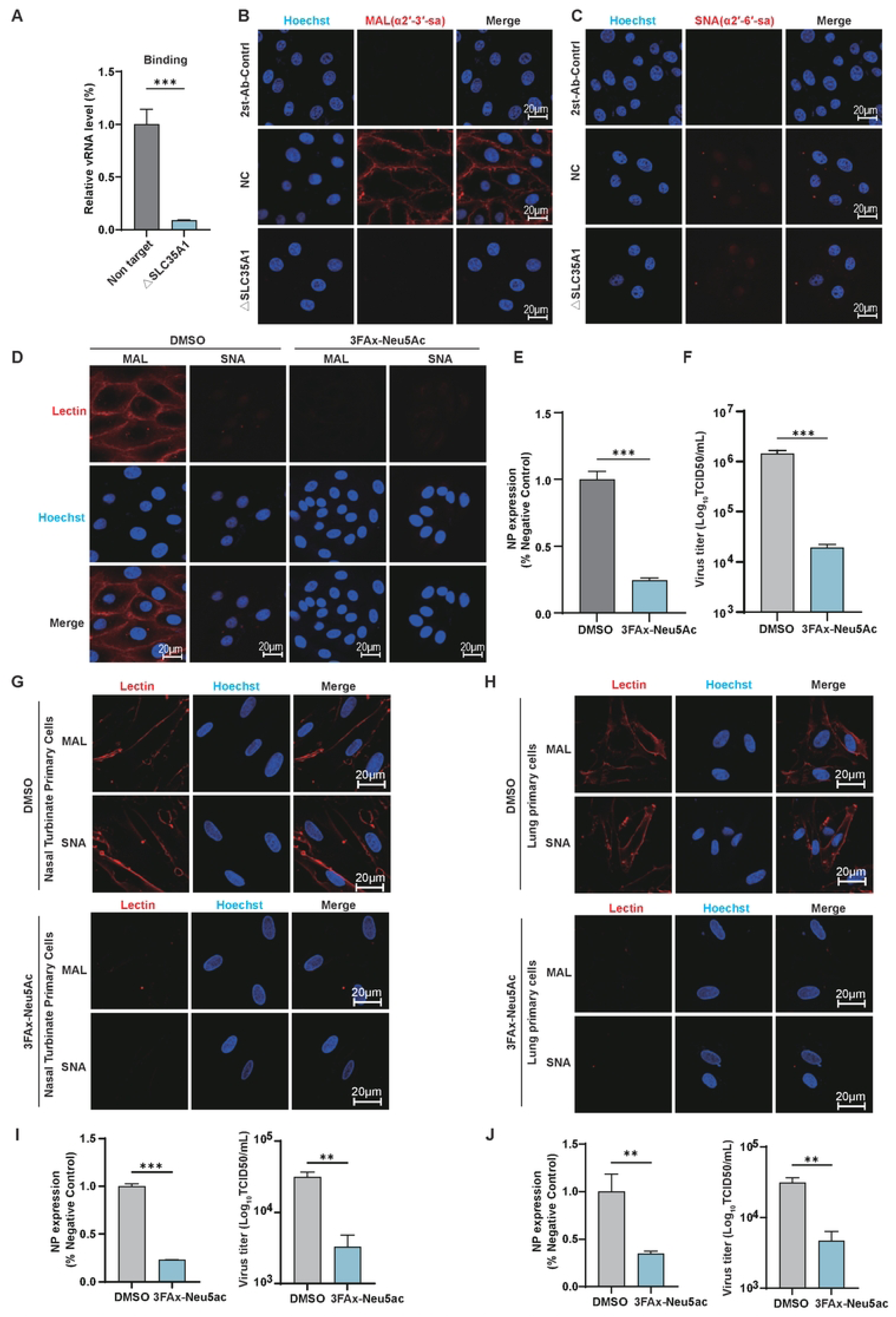
The sialic acid transporter SLC35A1 is essential for the binding of BPIV3 into MDBK cells. (A) Viral attachment was determined by qRT-PCR. Relative v-RNA level from three independent experiments with SD was statistically analyzed. (B) displays MDBK and SLC35A1 KO cells stained with α-2,3-linked sialic acid (SA), while (C) shows the same cell types stained with α-2,6-linked SA. The staining was performed using a second antibody control (2AbCtrl) to ensure specificity. The images were captured by confocal microscopy, and scale bars represent 10 μm. (D) Fluorescent microscopy of lectin binding in 3FAx-Neu5Ac-treated WT MDBK cells. SNA and MAL staining was performed as described for (B and C).MDBK cells were treated with either DMSO or 3FAx-Neu5Ac and subsequently infected with BPIV3 at a MOI of 1. After 24 hours, (E) the levels of viral nucleoprotein (NP) replication within the cells were assessed as a measure of infection. (F) The supernatant viral titers were also determined to evaluate the extent of viral replication and release. (G and H) Confocal microscopy was employed to examine the distribution levels of α-2,3- and α-2,6-linked sialic acid (SA) in bovine primary nasal turbinate cells and pulmonary fibroblasts, as well as in these cells treated with 3FAx-Neu5Ac. Furthermore, (I and J)bovine primary nasal turbinate cells and pulmonary fibroblasts treated with DMSO or 3FAx-Neu5Ac were infected with BPIV3 at a MOI of 1 for 24 hours. Subsequently, the viral titers in the cell supernatants and the expression levels of the nucleoprotein (NP) within the cells were determined.**,p < 0.002,***,p < 0.001.

Since α2,6-sialic acid was barely detected in MDBK cells, we sought to determine the expression of sialic acid in bovine respiratory cells. To this end, we isolated primary bovine nasal turbinate cells and pulmonary fibroblasts. These cells were treated with 3Fax-Neu5Ac to investigate the distribution of sialic acids and assess their impact on BPIV3 infection. Our results showed that both α2,3- and α2,6-linked sialic acids are present in these primary cells (Fig. 3G, H). A significant reduction in cell surface sialic acid levels was observed following treatment with the 3Fax-Neu5Ac inhibitor (Fig. 3G, H). We compared BPIV3 infection in cells treated to remove surface sialic acids with those treated with DMSO. Our results showed that BPIV3 infection was significantly reduced in cells lacking surface sialic acids (Fig. 3I, 3J). Collectively, these findings indicate that SLC35A1 facilitates the incorporation of sialic acid moieties onto cell-surface proteins, thereby serving as an essential host factor for BPIV3 binding.

### 2.4 LSM12 is involved in the entry of BPIV3 by affecting lysosomal acidification

Upon binding of BPIV3 to the cell surface, the virus fully exploits host endocytic pathways, including viral internalization at the cell surface, endosomal trafficking, and membrane fusion at the periphery of the nucleus[26, 27]. To elucidate how LSM12 deficiency confers resistance to BPIV3 infection, we first identified which steps of the BPIV3 life cycle are affected by LSM12. To assess its potential role in the entry of BPIV3, we first examined whether LSM12 is involved in viral attachment to the cells by quantifying viral nucleoprotein (NP) RNA (NP vRNA) levels of BPIV3 in cells deficient in LSM12 using qRT-PCR assay. The results revealed that there is no significant difference in NP vRNA levels between LSM12 KO cells and non-target cells (Fig. 4A). This indicates that LSM12 is not involved in facilitating the attachment of BPIV3. We assessed the potential role of LSM12 in subsequent entry stages of BPIV3. In our internalization assay, our results showed that there was a significant reduction in NP vRNA in LSM12 KO cells compared to non-target control cells (Fig. 4A). To confirm this observation, we analyzed viral transcription and genome replication in LSM12 KO cells using qRT-PCR. At 3 hours post-infection, NP vRNA levels were reduced by nearly 30% in LSM12 KO cells, suggesting a lower internalization of viral ribonucleoprotein (vRNP) (Fig. 4B).

**Fig 4.**
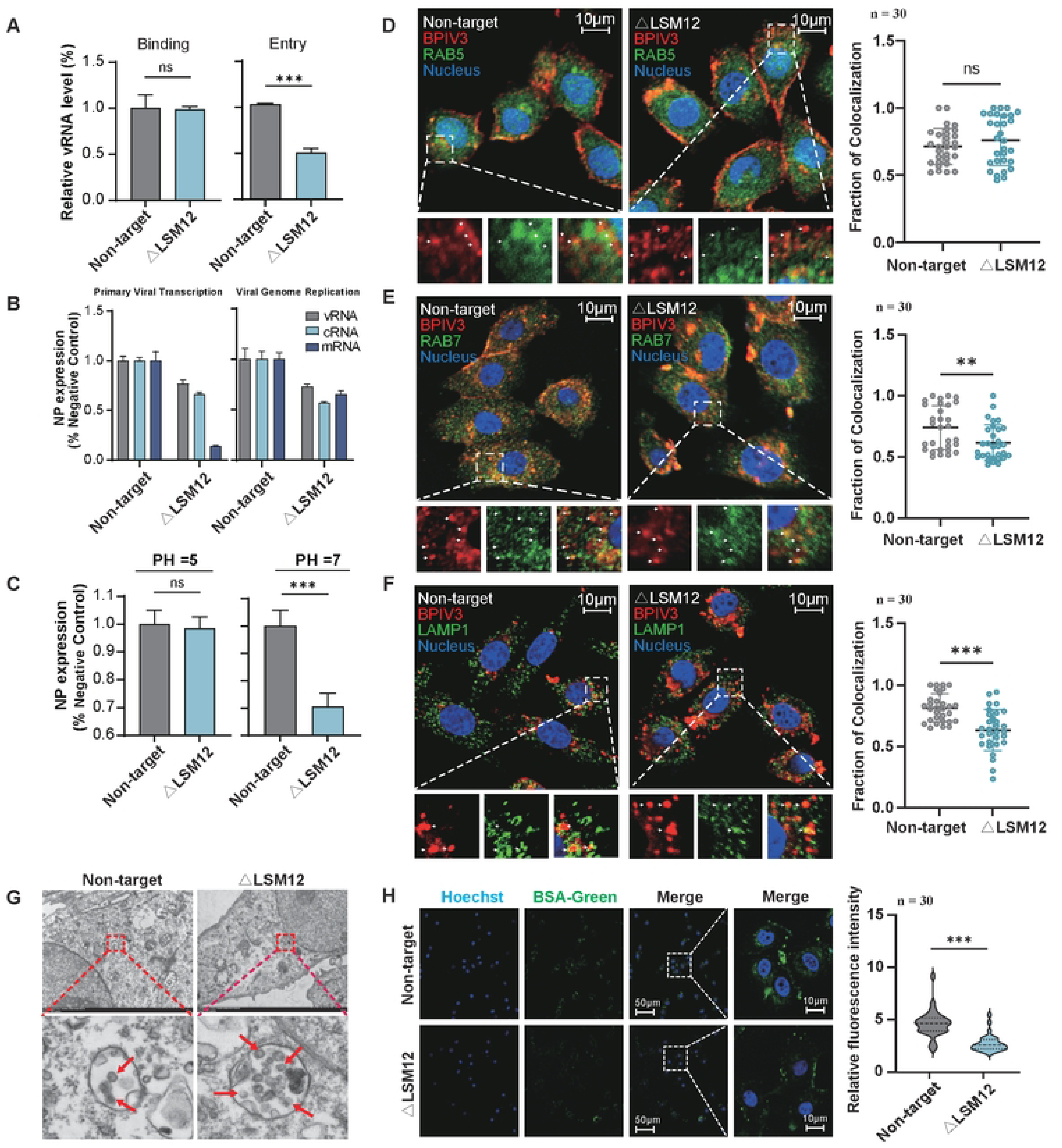
Knockout of LSM12 impedes the entry of BPIV3 by influencing lysosomal acidification. (A) Quantification of HN binding and internalization. The LSM12 gene knockout and non-target control cell lines were infected with BPIV3 (MOI = 1), and then the virus binding (A) on the cell surface and virus internalization. (B) qRT-PCR analysis of primary viral transcription and viral genome replication. Vector control and clonal KOs were infected with BPIV3 (MOI = 1) andviral NP vRNA, mRNA and cRNA levels were analyzed at 3 hpi or at 6 hpi. Data are represented as a percentage of expression relative to BPIV3-infected non-target cells ± SD. (C) Quantification of BPIV3-fused uncoating in LSM12-KO and non-target cells. (D-F) Colocalization of BPIV3 virus with endosomes and lysosomes was analyzed in non-target and LSM12-KO cell lines at different time points post-infection using confocal microscopy. At 10 minutes post-infection (D), colocalization with Rab5 (early endosomes) was assessed. At 20 minutes post-infection (E), colocalization with Rab7 (late endosomes) was evaluated. At 30 minutes post-infection (E), colocalization with LAMP1 (lysosomes) was examined. The following markers were used: Blue, Hoechst 33258 (nuclear stain); Red, R18-labeled BPIV3 virus; Green, GFP-tagged endosomal/lysosomal markers (Rab5, Rab7, LAMP1). Scatter plots (D-F) represent colocalization analysis of Rab5-positive, Rab7-positive, and LAMP1-positive structures with BPIV3 virus. Colocalization was quantified using the Coloc 2 plugin in Fiji (Mander’s colocalization coefficients). Data are presented as means ± SDs. Statistical significance was determined using a two-tailed Student’s t-test. For each group, 30 regions of interest were chosen within 3–5 cells each group. (G) Electron microscopy images of BPIV3 virus infection at 40 min post-infection in non-target MDBK, LSM12 KO cells. (H) Confocal microscopy analysis of LSM12-KO or non-targeting sgRNA using DQ-green bovine serum albumin (BSA). Cells were incubated with 20 µg/mL DQ-BSA. The fluorescence intensity of DQ-BSA was quantified by dividing the total fluorescence intensity by the number of cells within each frame. The violin plot depicts the relative fluorescence intensity for different sgRNAs. Scale bar = 20 µm. Error bars represent the standard deviation from three randomly chosen frames.***,p < 0.001,ns, non-significant.

This is likely due to inefficient release of vRNPs into the perinuclear space as a result of impaired endocytic pathways. At 6 hours post-infection, the reduction in NP vRNA reached 40%, with comparable decreases in viral NP complementary RNA(cRNA) and viral NP transcript (NP mRNA) mRNA observed in LSM12 KO cells relative to control cells (Fig. 4B). This observation suggests that the initial transcriptional defect impaired subsequent genome replication.

Till now, our findings have established the critical role of LSM12 in BPIV3 infection. Next, we further interrogate the potential role of LSM12 in BPIV3 fusion. Given its molecular property, we expect that LSM12 is likely involved in regulating pH change required for the BPIV3 fusion process. To test this hypothesis, we performed the “acid bypass” assay [28]. In this assay, viral fusion was triggered at the plasma membrane by acid-buffer treatment to circumvent internalization defects. Our results showed that unlike the control groups, which showed no change in NP vRNA levels after acid treatment, LSM12 KO cells showed an upregulation of NP vRNA levels in a single round of infection after acid treatment (Fig. 4C). This suggests that pH treatment rescued the deficiency of LSM12, which implies that LSM12 is actively involved in facilitating BPIV3 fusion via regulating pH.

To confirm that LSM12 plays a critical role in membrane fusion during the BPIV3 entry process, we tracked BPIV3 cellular localization using our purified BPIV3 particles labelled with lipophilic octadecyl rhodamine B chloride (R18) probes (BPIV3-R18) [29, 30] via confocal microscopy analysis. In both non-target and LSM12 KO cells, the BPIV3-R18 showed no difference in colocalization with Rab5 endosomes (early endosome) at 10 minutes post-infection (Fig. 4D). However, at 20 minutes post-infection, the internalized virus exhibited significantly lower colocalization with Rab7 endosomes (late endosome) in LSM12 KO cells than in non-target cells (Fig. 4E). At 30 minutes post-infection, BPIV3-R18 levels, which partially colocalized with Lamp1 (lysosome), were significantly decreased in LSM12 KO cells compared to non-target cells (Fig. 4F). These results suggest that LSM12 deficiency does not perturb early endosomal (Rab5) trafficking but impairs subsequent late endosomal (Rab7) maturation and lysosomal (Lamp1) targeting, thereby blocking viral progression through the endosomal-lysosomal axis.

Moreover, we also examine the trafficking and degradation of incoming BPIV3 virions within endo-lysosomal compartments. We used electron microscopy to analyse infected LSM12 KO cells alongside non-target cells at an MOI of 20, after 40 minutes of infection. Our findings revealed that, unlike control cells, internalized viral particles were significantly concentrated within endosomal vesicles in LSM12 KO cells (Fig. 4G). The results imply that the fusion process between the viral membrane and the endosomal membrane is impaired in LSM12 deficient cells.

To confirm that LSM12 is involved in lysosomal acidification, we incubated LSM12 KO and non-target cells with DQ-Green BSA. Our results showed that lysosomal degradation of DQ-Green BSA was only suppressed in LSM12 KO cells compared to non-target cells, which strengthened the involvement of LSM12 in lysosomal acidification (Fig. 4H).

To further verify the influence of LSM12 in BPIV3 infection, the corresponding gene complement cell lines were generated using a retrovirus-induced stable expression system with a same-sense mutation at the sgRNA-targeted PAM region (Fig. 5A). Subsequently, knockout and LSM12 complement cell(C-ΔLSM12) lines were infected with BPIV3 at an MOI of 0.1. Viral titers and RT-PCR data showed that LSM12 over expression in knockout cells increased BPIV3 replication compared to control cells (Fig. 5B). Confocal microscopy results revealed that the colocalization of BPIV3-R18 with LAMP1 at 30 minutes post-infection was not significantly different between the non-target and C-ΔLSM12 groups (Fig. 5C). This suggests that the viral membrane fusion process, which depends on lysosomal engagement, was effectively restored in the complemented C-Δ LSM12 cells. The DQ-Green BSA assay confirmed a restoration of lysosomal acidification in C-ΔLSM12 cells compared with the levels observed in LSM12 KO and negative control (NC) cells (Fig. 5D). In conclusion, these results demonstrated that LSM12 affects membrane fusion of BPIV3 by influencing the reduction of lysosomal pH.

**Fig 5.**
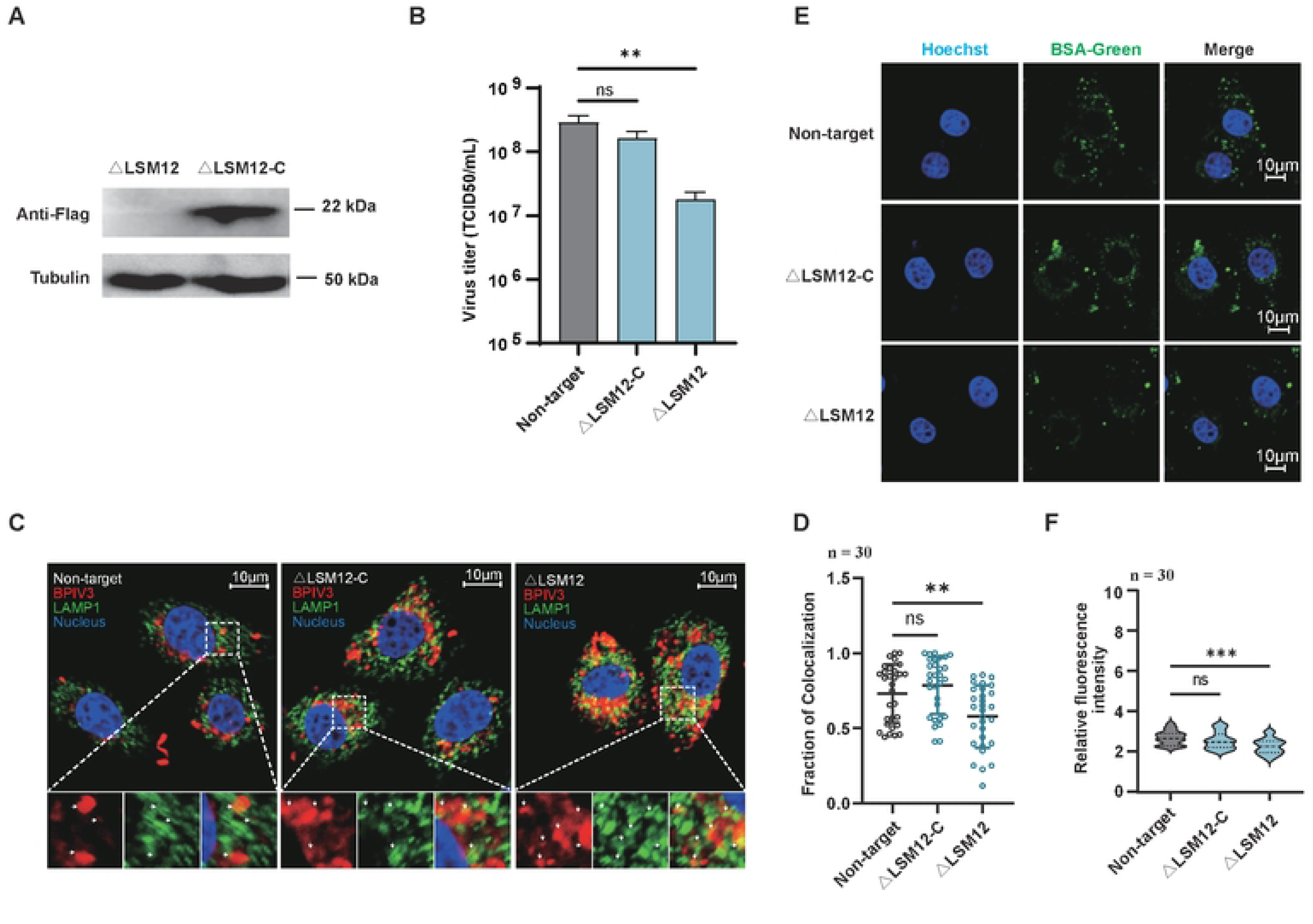
Overexpression of LSM12 restores the entry of BPIV3 into the LSM12 KO cells. (A) Western blot analysis of retrovirus-induced LSM12 genes complement expression. (B) LSM12 knockout and complement (FLAG-tagged) cell lines were infected with BPIV3 (MOI = 0.01) and viral titers were measured at the indicated times on MDBK cells. (C) Colocalization analysis of BPIV3 and lysosomes in non-target, LSM12 knockout, and complement cell lines at 30 min by confocal microscopy. Lysosomes were labeled with FITC-conjugated anti-LAMP1 antibody. Blue, Hoechst 33258 (nuclei); red, R18-labeled BPIV3; green, FITC-labeled LAMP1 (lysosomes). Scale bar = 10 µm. Scatter plot (D) represents colocalization analysis of (C). Colocalization analyses are consistent with the previous description. (E) Confocal microscopy analysis of non-targeting, LSM12 knockout and complement cells using DQ-green bovine serum albumin (BSA),Scale bar = 10 µm. Violin plot (F) depicts the relative fluorescence intensity for different cells, colocalization analyses are consistent with the previous description.Scale bar = 10 µm.

### 2.5 LSM12 affects lysosomal acidification through TPC channels

Given that NAADP is a potent Ca2+ mobilizing intracellular messenger that induces Ca2+ release from lysosome-like acidic organelles and LSM12 binds directly to NAADP via its LSM domain, it is reasonable to hypothesize that this interaction could result in NAADP-evoked TPC activation and Ca2+ mobilization influencing acid stores [31, 32]. Two important signaling molecules control the opening of TPCs: PtdIns(3,5)P2, an inositide generated from the phosphorylation of PtdIns(3)P by the five kinase PIKfyve, which also activates the TRPLM1 channel, and NAADP [33, 34] (Fig. 6A). We expected that these two factors would also be required for BPIV3 infection. To this end, we used the specific drug target for each individual factor and tested viral infection in these treatments. We first used a known PIKfyve suppressor, apilimod. Our entry analysis with BPIV3 pseudovirions revealed that there is reduced entry of BPIV3 pseudovirions into MDBK cells in a dose-dependent manner (Fig. 6B). We next treated cells with Ned19, a known NAADP antagonist, and observed similar reduction. The TRPML1 inhibitor (1R,2R)-ML-SI3 did not influence viral replication, as the decrease in cell viability was significantly lower than the decrease in viral titer (Fig. 6B). To test whether the inhibitor would inhibit BPIV3 entry, we used confocal microscopy to study the colocalization between BPIV3-R18 and anti-Lamp1 (lysosomes) after treating cells with the inhibitors for 1 hour. Treatment with Ned19 decreased viral entry, while blocking TPC and TRPML1 activity with apilimod, an inhibitor of TPC and TRPML1, decreased BPIV3 entry (Fig. 6C). However, treatment of cells with (1R,2R)-ML-SI3, a TRPML1 inhibitor, had no effect (Fig. 6C, D), indicating that TPC, and not TRPML1, is important for BPIV3 entry. We next asked whether the three inhibitors affected lysosomal function, using the DQ-BS probe. As expected, (1R,2R)-ML-SI3 had no effect, but treatment with Ned19 and apilimod significantly reduced the fluorescence of DQ-BSA compared to non-target cells(Fig. 6E, F).

**Fig 6.**
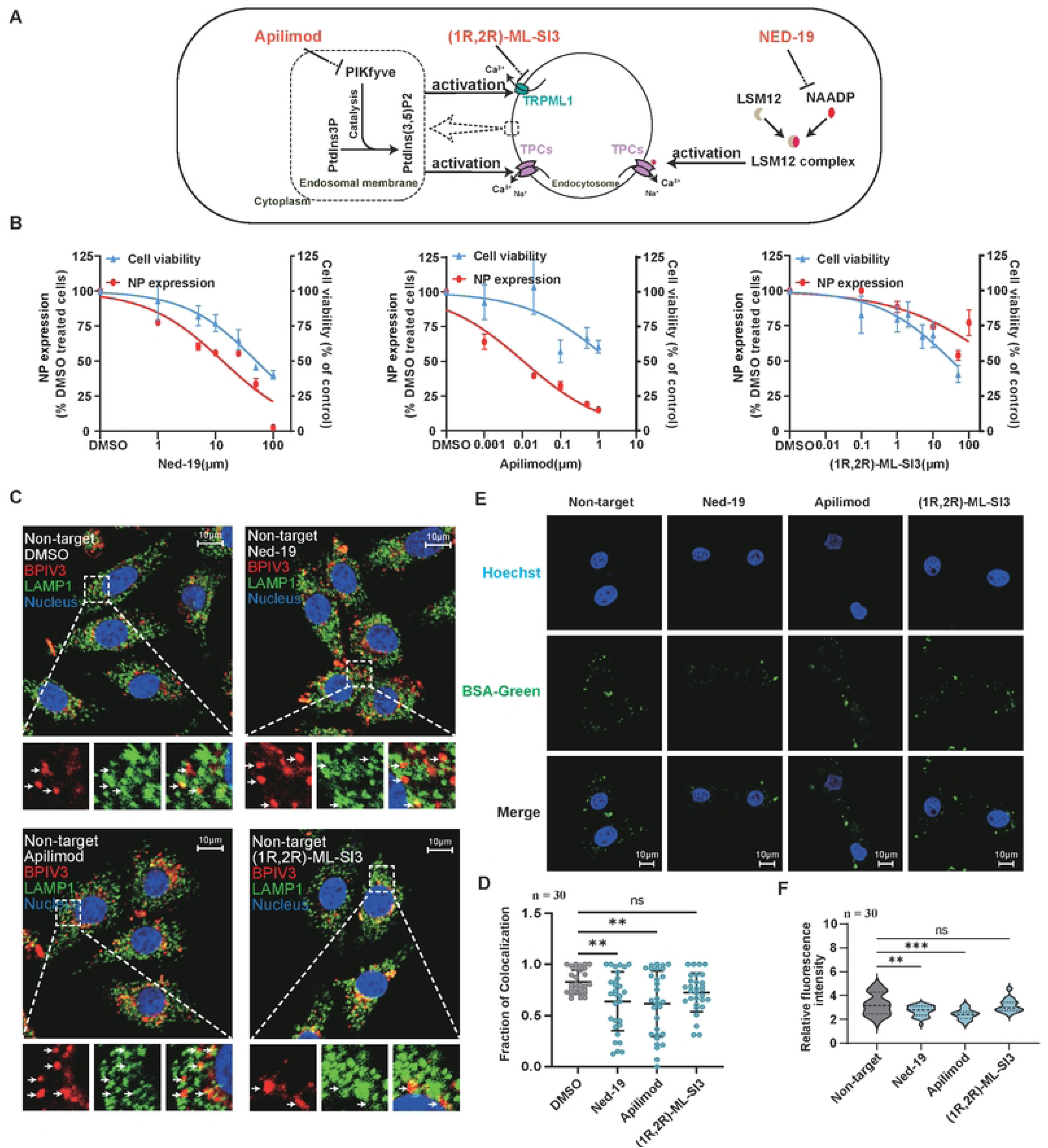
LSM12 influences the entry of BPIV3 by activating the TPC channel of lysosome. (A) Inhibition of Calcium Signaling Pathways by Apilimod, (1R,2R)-ML-SI3, and NED-19 in Endosomal Compartments. (B) Cell viability (blue) and NP expression (red) were evaluated in cells treated with escalating concentrations of Ned-19 (left panel), Apilimod (middle panel), and (1R,2R)-ML-SI3 (right panel). Cells were pre-treated with the compounds for 30 minutes, followed by infection with BPIV3 at MOI of 0.01. After a 24-hour infection period, viral infection was quantified using qPCR, and cell viability was assessed with the CCK-8 assay. Data are presented as the mean ± SD of triplicate measurements from a single experiment, with consistent results observed in at least three independent replicates. (C) Confocal microscopy was employed to evaluate the colocalization of BPIV3 with lysosomes in cells pretreated with Apilimod (0.01µM), (1R,2R)-ML-SI3 (0.1µM), and NED-19 (10µM) for 30 minutes. Lysosomes were labeled with FITC-conjugated anti-LAMP1 antibody. Blue, Hoechst 33258 (nuclei); red, R18-labeled BPIV3; green, FITC-labeled LAMP1 (lysosomes). Scale bar = 10 µm. (D) Scatter plot representing the colocalization analysis of panel (C). Colocalization analyses are consistent with the previous description.n=30. (E) Confocal microscopy analysis of endocytosis in Apilimod (0.01µM), (1R,2R)-ML-SI3 (0.1µM), and NED-19 (10µM) pretreated MDBK cell lines using DQ-green bovine serum albumin (BSA). Scale bar = 10 µm. (F) Violin plot depicting the relative fluorescence intensity for different cell lines, which corresponds to the endocytosis levels shown in panel (E). Colocalization analyses are consistent with the previous description. n=30.

### 2.6 Ned19 inhibits NAADP-LSM12-TPC signaling axis to restrict viral infection

Inspired by emerging evidence linking NAADP signaling to viral entry and replication in viruses, we investigated whether pharmacological inhibition of NAADP with Ned19 could broadly restrict viral infection. In this study, we treated cells with Ned19 and infected them with viruses including ORFV, BoHV-1, VN/H5N1, COW/H5N1 and HSV, while using DMSO treated cells as controls. Virus titers were determined by standard plaque assays or TCID_50_ methods, and cell viability assays were performed to evaluate the toxicity of Ned19. Results showed that viral titers in Ned19-treated cells were significantly lower than those in DMSO treated cells for all tested viruses (P<0.02) (Fig. 7A-E). Meanwhile, Ned19 showed relatively low cytotoxicity at the effective concentration of 10 μM/mL, with cell viability maintained above 80% (Fig. S3). These findings demonstrate that Ned19 effectively inhibits viral infection by suppressing the NAADP-LSM12-TPC signaling axis, thereby exhibiting promising broad-spectrum therapeutic potential against novel emerging infectious diseases lacking effective treatment options.

**Fig 7.**
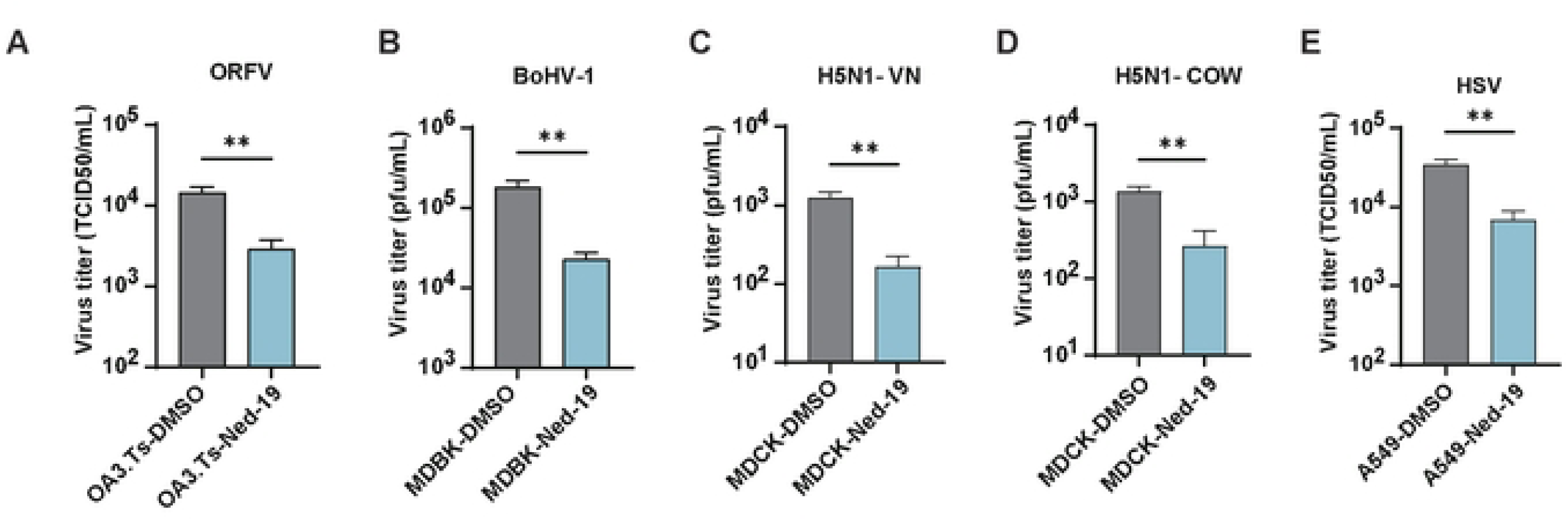
Ned19 suppresses replication of multiple viruses. (A) OA3.Ts cells (ORFV), MDBK cells (BoHV-1), MDCK cells (H5N1-VN and H5N1-COW), and A549 cells (HSV) were pre-treated with 10 µM/mL Ned19 for 1 hour, followed by viral infection at an MOI of 0.01 for 1 hour. Subsequently, cells were incubated at 37℃ for an additional 24 hours. Supernatants were collected and virus titers were determined by TCID50 or plaque assay. Data are presented as the mean ± SD (n = 3). **p < 0.002.

Overall, our study presents the establishment of BovGeCKO, a genome-wide CRISPR knockout library for bovine cells, and its application in identifying host factors essential for BPIV3 infection. The key genes identified include SLC35A1, required for viral attachment via sialylation, and LSM12, which facilitates viral entry through lysosomal acidification via TPC channels. These findings reveal critical host-virus interaction mechanisms and fill the gap in understanding the entry of BPIV3.

## 3 Conclusion

Our findings demonstrate the establishment of an efficient CRISPR/Cas9-based screening for functional analyses in bovine cells, which was further validated by our results that identified key host factors for BPIV3 entry. As described in our preliminary model (Fig. 8), BPIV3 binding and entry involve factors such as the sialic acid transporter SLC35A1, which was highly enriched in our screen due to its necessity for the viral receptor and HN binding. Additionally, we found that LSM12 is critical for BPIV3 endocytosis. Loss of LSM12 inhibits viral fusion by disrupting lysosomal acidification, mediated by TPC ion channel modulation.

**Fig 8.**
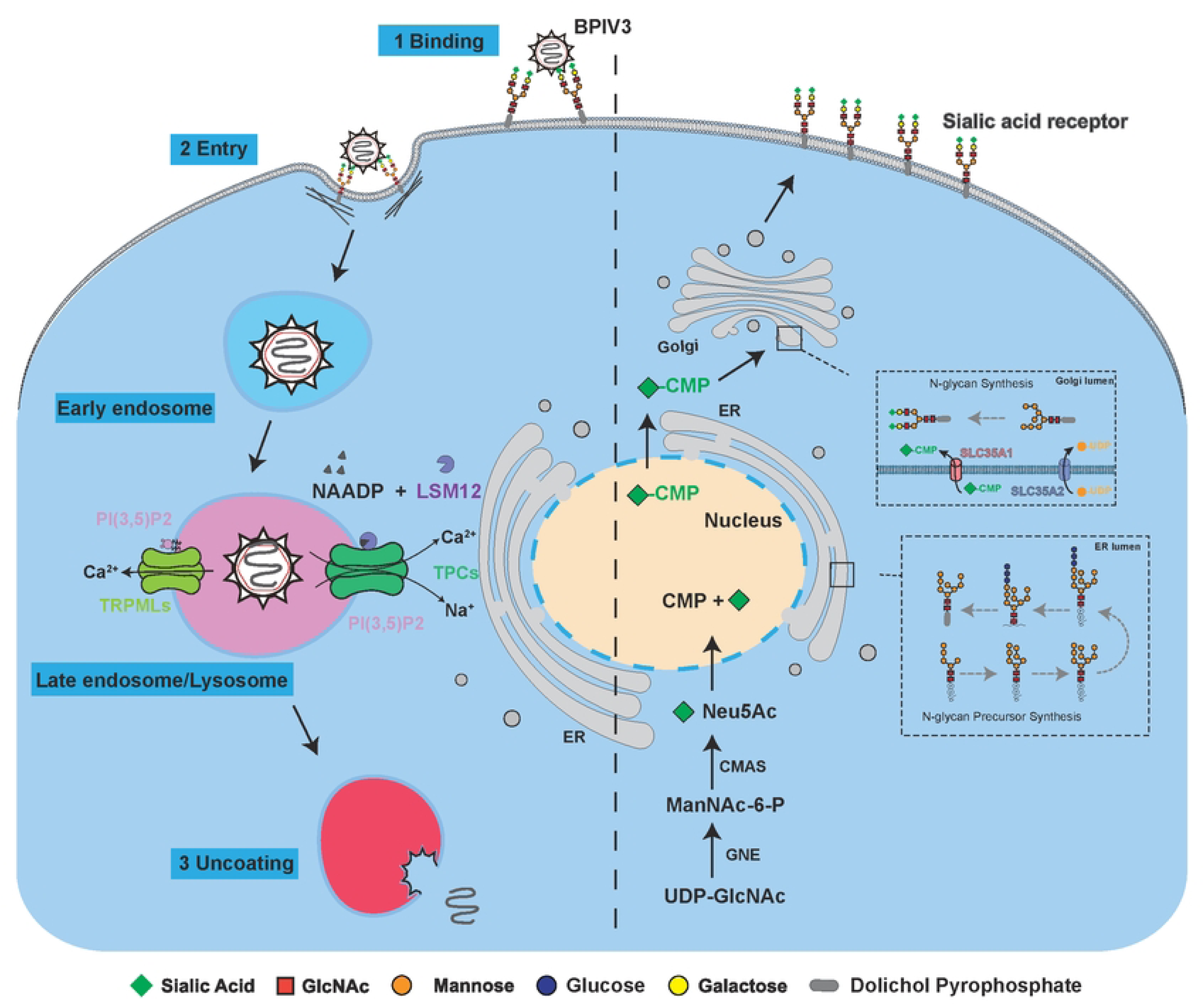
Model of BPIV3 Entry into Bovine Cells. This diagram illustrates the sequential steps involved in the entry and intracellular trafficking of BPIV3 virus within host cells. The process begins with the binding of BPIV3 to sialic acid receptors on the cell surface (Step 1). Following binding, the virus is internalized via endocytosis (Step 2), leading to its enclosure within early endosomes.The virus then traffics to the late endosome/lysosome, where it undergoes further maturation. A complex consisting of LSM12 and NAADP can activate TPCs channels. This activation triggers the release of calcium (Ca²⁺) and sodium (Na⁺) ions from the late endosome/lysosome, resulting in lysosomal acidification that facilitates virus membrane fusion and entry into the cytoplasm (Step 3). The diagram also highlights the synthesis and trafficking of CMP-Neu5Ac, a key component in sialic acid receptor biosynthesis. CMP-Neu5Ac is generated in the nucleus from UDP-GlcNAc and transported to the Golgi apparatus, where it is utilized in the synthesis of sialic acid receptors on the cell surface. This process is essential for viral recognition and entry. Overall, this schematic provides a comprehensive overview of the molecular mechanisms underlying BPIV3 virus entry and intracellular trafficking, emphasizing the critical roles of various cellular components and pathways in the viral life cycle.

In constructing this cellular library, we took into account the potential validation limitations of gene editing operations (modification of TRIM5α) observed in previous studies (Tan et al., 2023). To eliminate phenotypic confusion that genetic manipulation might introduce, we chose to construct the library using MDBK cells that had not been genetically modified. By building upon and optimizing the previously used method, our system can directly capture the dynamic interaction network between viruses and the entire host genome in its natural state. Furthermore, in pooled CRISPR screens, excessive sgRNA delivery efficiency (greater than 60%) poses a risk of introducing multiple sgRNAs per cell, which may lead to nonspecific toxicity [35, 36]. Adhering to Poisson distribution principles, maintaining infection rates between 20% and 60% optimizes the prevalence of single-integrant cells while reducing both uninfected cells and the initial cell population requirements [36]. Our CRISPR libraries use a low MOI (MOI=0.3) and a high coverage depth (1000×) to ensure the integrity and validity of the library. Additionally, we conducted multiple rounds of BPIV3 lethal infection to enrich for key host genes. Deep sequencing revealed that while 613 gRNAs were absent from the CRISPR knockout cell library we constructed, only 23 genes were lacking. Our study further demonstrated that BPIV3 replication was significantly reduced in both SLC35A1 and LSM12 knockout cell lines compared to non-target cells. Therefore, the GeCKO screening resource and the high-stringency screening strategy developed in this study are robust and reliable.

SLC35A1, a member of the solute carrier family 35 responsible for transporting CMP-sialic acid into the Golgi apparatus, plays an essential role in cell surface sialylation [37, 38]. While its importance has been revealed in the context of other viral infections, such as influenza A virus where sialic acid acts as a key receptor [15, 39, 40], the specific involvement of SLC35A1 in BPIV3 infection had not been established. Our study now provides the first evidence linking SLC35A1 to BPIV3 infection. In our GeCKO screen, SLC35A1 surfaced as the most significantly enriched hit, with gene knockout resulting in a four-log decrease in BPIV3 infection. This indicates the critical role of SLC35A1, likely by providing the necessary sialic acid ligands for viral attachment to the host cell surface. We further examined the sialic acid profiles on the surface of our working MDBK cells and found an absence of α2,6-linked sialic acids. Treating primary cells with 3F-Neu5Ac, an inhibitor targeting downstream sialyltransferases, significantly reduced BPIV3 infection. These observations suggest that BPIV3 preferentially uses α2,3-linked sialic acids for viral attachment, emphasizing the broader importance of cell surface glycosylation in BPIV3 infection and indicating that modulating sialic acid levels could serve as a potential antiviral strategy.

Another factor that we further characterized is LSM12, which belongs to the LSM family and is reported as a selective, high-affinity receptor for NAADP, mediating its association with TPCs and subsequent Ca²⁺ mobilization from acidic stores [31]. While its role in calcium homeostasis is established [31], the involvement of LSM12 in viral infection, particularly BPIV3, remained poorly understood. Our study provided direct evidence to show a critical role for LSM12 in BPIV3 infection. We found that LSM12 restricts viral internalization. Based on this observation, our further mechanistic investigations revealed the importance of intracellular Ca²⁺ homeostasis for viral internalization. Additional experiments showed that viral fusion is blocked in LSM12 KO cells. Instead of progressing normally to late endosomes (positive for Rab7), virions accumulated within lysosomes (positive for LAMP1) at later time points, a phenotype supported by decreased endocytic cargo degradation in KO cells. This suggests a failure of pH-dependent fusion within the endo-lysosomal pathway due to impaired endosomal acidification. Therefore, our results indicated that LSM12 is involved in BPIV3 internalization, and its knockout impairs viral fusion by affecting endosomal acidification.

In summary, our study used CRISPR/Cas9 screening to interrogate the mechanisms of BPIV3 infection in bovine cells. We identified two critical host factors: the sialic acid transporter SLC35A1, essential for viral receptor expression and initial viral binding, and the protein LSM12, which is crucial for viral endocytosis. We demonstrated that LSM12 knockout significantly impairs viral fusion by disrupting endosomal acidification. These findings not only advance our understanding of BPIV3 entry but suggest new avenues for antiviral development.

## Acknowledgements

We would like to express our sincere gratitude to Dr. Xin Huang for the isolation of BPIV3 strain XJA13, which provided crucial materials for this study. We also thank Dr. Jie Gao for kindly providing the BoHV-1 and HSV-1 F strain, which were essential for our experimental work. Additionally, we appreciate the efforts of the members of our laboratory who maintained the Orf virus (ORFV), A/Viet Nam/1203/2004 H5N1 (VN/H5N1), DC/H5N1, and the PR8-derived H5N1 recombinant virus COW/H5N1, ensuring the smooth progress of our research.

## Author contributions

G.J.W., W.H., and R.H. conceived and designed the experiments; Y.J.H., X.R.G., Z.Y.S., X.G., Y.S., Y.T., and S.L. performed the experiments; G.J.W., Y.J.H., and X.R.G. analyzed the data; J.G., X.H., and Q.Y.Z. provided the viruses. G.J.W., J.L.W., Z.Z., and R.H. interpreted the data; Y.J.H., R.H., and G.J.W. wrote the original draft; G.J.W., C.L., and R.H. reviewed and edited the paper.

## Supplemental information

**Fig S1. Generation of 34 MDBK-Cas9 monoclonal cells and depiction of individual sgRNA enrichment for selected hits. Related to Figure 1**

(A) MDBK cells were seeded in 6-well plates. The next day, the medium was replaced with 2 mL of culture medium containing 200, 100, 50, 20, 10, or 0 μL of lentivirus supernatant mixed with 8μg/mL polybrene. After 48 hours, cells were collected for FACS analysis to determine the percentage of GFP-positive cells. The virus amount used for wells with ≤30% transduced cells was selected to calculate the virus amount required for screening at MOI=0.3.

(B) Cutting efficiency of M-Cas9-clones in MDBK cells was quantified by TIDE analysis. An sgRNA targeting CPT1B was randomly selected to evaluate the cleavage activity of the Cas9 protein expressed in the M-Cas9 cells.

**Fig S2. Generation of seven candidate gene knockout monoclonal cells and sgRNA lentivirus titration. Related to Figure 2**

(A) A 500 bp to 800 bp region flanking the sgRNA target site was amplified in WT and sgRNA knockout monoclonal cells. The primers used are shown in Table S3. Yellow letters, PAM region; red region, nucleotide mutations; Blue region, sgRNA target sequences. Related to Figure 2A.

(B) Normalized read counts for individual sgRNAs compared between round1, round2, round3 and cell control. The four or five sgRNAs are represented by different dotted lines.

**Fig S3. Effect of Ned19 on cell viability. Related to Figure 7**

(A) Cell viability was assessed in cells treated with 10µM/mL Ned-19. After 24 hours of Ned-19 treatment, cell viability was assessed using the CCK-8 assay.

**Table S1. MAGeCK Analysis of Three-Round CRISPR Screens. Related to Figure 2**

This supplementary table (Table 1) presents data from three - round MAGeCK analysis performed in CRISPR screens, which is relevant to the host - gene screening process related to BPIV3 as detailed in our study.

**Table S2. MaGplotR Analysis of Three-Round CRISPR Screens. Related to Figure 2**

This supplementary table (Table S2) presents results from three-round MaGplotR analysis, which builds upon the MAGeCK analysis outcomes (detailed in Table S1) for further data interpretation and visualization.

**Table S3. Gene ontology analysis of 0.5% of the ranked hits from the result of the MaGplotR analysis. Related to Figure 2**

This supplementary table (Table 3) presents the Gene Ontology (GO) analysis of 0.5% of the ranked hits from the MaGplotR analysis results, highlighting enriched biological processes, molecular functions, or cellular components associated with the identified host genes in the context of BPIV3 screening.

**Table S4. Primers used in this research. Related to STAR Methods.**

This supplementary table (Table S4) lists the primers used in this research, corresponding to the details described in the Methods.

## 4 STAR Methods

### 4.1 Cell culture and virus

MDBK cells, HEK293T, MDCK, A549, OA3.Ts cells were maintained in our laboratory (ATCC source) and cultured in Dulbecco’s modified Eagle’s medium (DMEM, Thermofisher, #12100061) containing 10% fetal bovine serum (Cellsera. #F42006). Bovine lung and nasal progenitor cells were isolated from bovine embryos and cultured in DMEM medium containing 10% fetal bovine serum(D10) (Gibco, #10099141). BPIV3 strain XJA13 was isolated by Xin Huang [41]. Orf virus (ORFV), A/Viet Nam/1203/2004 H5N1 (VN/H5N1), DC/H5N1 were maintained in our laboratory. The COW/H5N1 was a PR8-derived H5N1 recombinant viruses, in which haemagglutinin (HA) and neuraminidase (NA) were originated from A/dairy cow/Texas/A240750066-18/2024(H5N1) and the internal genes were originated from A/PuertoRico/8/34 H1N1 (PR8). The mult-basic cleavage site of HA of VN/H5N1, and COW/H5N1 was removed to reduce virulence. BoHV-1,HSV-1 F strain were provided by Jie Gao.

### 4.2 Plasmids

pSPAX2 (Addgene#12260), pMD2.G (Addgene#12259), lenti-guide-puro (Addgene#52963), lenti-blast-Cas9 (Addgene#52962), and lenti-Cas9-gRNA-puro (Addgene#52961) were purchased from addgene. The pLV3-CMV-MCS-3×FLAG-Neo was constructed in our laboratory.

### 4.3 Viral production and infection

For lentivirus production, 2×10⁶ HEK293 cells were first distributed across a 6-well plate. To illustrate, an EP tube labelled as Tube A was initially removed, 150 μL of opti-MEM was added, along with 2.4 μL of PlusTM reagent, and the contents were mixed and left to stand. Another EP tube should be removed and labelled as B tube. Subsequently, 150 μL of opti-MEM and a total of 2.4 μg of plasmid should be added. The ratio of the three plasmids is (plenticas9-Blast/plenti-GFP/plenti-gRNA): pVSVg:pGagPol = 3:5:4. Finally, 7.5 μL of LTX transfection reagent should be added. The contents of the tube should be mixed thoroughly and incubated for three minutes. The tube A contents should then be mixed with tube B and left to stand at room temperature for 15 minutes. The cell plate should be removed, the AB mixture added to the medium after cell exchange, and the plate returned to the CO₂ incubator after slight shaking of the cross. Following a 48-hour incubation period, the supernatant containing the lentivirus was aspirated and 1 mL of D10 medium was added. Twenty-four hours later, the venom was collected once more and thoroughly combined. The collected lentiviral venom was subjected to centrifugation at 10,000 rpm for 10 minutes at 4°C. Following this, the centrifuged venom was filtered through a 0.45 μM filter, after which it was dispensed and stored at -80°C.

For the replication of BPIV3, negative control cells with CRISPR knockdown MDBK cells were plated at a density of 2 × 10⁶ cells per well in 6-well plates. The plate was aspirated in order to discard the old medium, and the cells were washed three times with phosphate-buffered saline (PBS,Thermofisher, #10010023). BPIV3 was then introduced to the cells at a multiplicity of infection of 0.1 for one hour at 37°C. The cells were subsequently washed three times with PBS, and 2mL of DMEM medium was added to the culture. Once the cell lesions reached 70% of their maximum size, the viral solution was collected and subjected to centrifugation at 13,000 rpm for 10 minutes at 4°C using a centrifuge. It was then filtered through a 0.45 μM filter and stored in a -80°C refrigerator.

### 4.4 Cow Genome-wide library design

The bovine genome-wide information reference sequences, Bos taurus UMD 3.1.1 and Btau 5.0.1, were downloaded from the NCBI database in August 2022.Firstly, the sequences were indexed using Bowtie and the CDS sequences of all genes were extracted. Subsequently, CRISPRCasFinder (v3.1.0) was employed to identify gRNAs with a mismatch of at least three bases. Ontarget scores were then calculated and gRNAs with high scores were selected [18]. For genes with an inadequate design, CFD scores were calculated for the second round of selection.

### 4.5 Construction of a genome-wide sgRNA library plasmid

The gRNA sequences were commissioned to be synthesised by Nanjing Genscript Co. For library amplification, the library plasmid and Stbl4 sensory fine (Invitrogen#11635018) were gently mixed according to 1 μg of library plasmid with 200 μL of sensory cells, respectively, using an electrotransfer cup (BIO-RAD#1652083), and added to the bottom of the electrotransfer cup according to 20 μL as a unit and inserted on ice. Electrotransformation (1.2 KV, 25 μV, 200 Ω) was carried out using a Burroughs Electrotransferometer (BIO-RAD#165-2660), and at the end of the electroshock pulse, 1 mL of SOC (Invitrogen#15544034) medium was quickly added to the electrotransfer cup, and the sensory cells in the electrotransfer cup were gently blown with a pipette gun, left to mix, and then transferred to 15 mL shaker tubes. After repeating the above steps for 10 times, the shaking tube was put into a shaker at 37 °C, 225 rpm/min, shaking the bacteria for 1 h, and LB plate was applied. 16 h later, the total number of plaques was counted, and all plaques were collected only if the total number of plaques was greater than 200, and plasmids were extracted using the HiSpeed Plasmid Maxi Kit (QIAGEN#12663) according to the manufacturer’s requirements. Plasmid libraries were amplified by PCR using Takara Ex Taq (RR001Q) (28 cycles), and the products were purified using FastPure Gel DNA Extraction Mini Kit (Vazyme#DC301-01) before being tested for library coverage by high-throughput sequencing. Amplification primer reference table S4.

### 4.6 Generation of Cas9-expression cell line

The MDBK cell culture medium was replaced with DMEM containing 8 μg/mL polybrene, 10% FBS, and the Cas9-Puro lentivirus was introduced to the medium at an multiplicity of infection (MOI) of 0.3. Following a 48-hour incubation period, the medium was replaced with DMEM containing 2.2 μg/mL puromycin and 10% FBS. Subsequently, after a five-day period, the resistant cells were harvested, and 32 monoclonal cells were obtained through the limited cell dilution method. The monoclonal cells that stably expressed the Cas9 gene and exhibited the highest knockdown efficiency were obtained through Western blot and knockdown efficiency detection (TIDE) [17].

### 4.7 Gene editing efficiency testing by T7E1 digestion and TIDE analysis

Seven days after transfection with individual CPT1B sgRNA, crude genomic DNA was extracted from cells using Quick-DNA Miniprep Plus Kit (ZYMO RESEARCH #D4068) following the product manual. The extract was diluted 1:10 in nuclease free water and 1ul of the dilution used for PCR using primers flanking the target site and PrimeSTAR® GXL DNA Polymerase with supplied buffer, producing amplicons between 300-800bp in size. 2ul of PCR reaction were run on 2% Agarose gel to estimate amplicon concentration. Without purification the calculated volume of PCR reaction containing ∼200ng PCR product based on the agarose gel was denatured at 95°°C for 5 minutes and re-annealed by dropping the temperature to 25°C°C at 0.1°C per second in a thermal cycler, at the end of the program the temperature was reduced to 4°C. Right after re-annealing, 1ul T7 Endonuclease I (New England Biolabs#M0302S)was added to the PCR reaction directly and incubated at 37°C for 30 minutes. Immediately after incubation, the reaction was analyzed on a 2% Agarose gel. The editing efficiency was also examined by TIDE analysis (https://tide.deskgen.com/) following Sanger sequencing of purified PCR products.

### 4.8 Lentivirus titration

MDBK cells were seeded in 6-well plates at a density of 2x10^5^ cells per well using 2 mL culture medium consisting of DMEM, 10% fetal bovine serum (FBS) and no antibiotics. The cells were cultured overnight to reach 30% to 50% confluence. The following day, the medium was replaced with 200ul, 100ul, 50ul, 20ul, 10ul or 0ul of lentivirus supernatant mixed separately with 8μg/mL polybrene in culture media with a total volume of 2mL. After 24 hours of incubation, the virus was removed and 2mL of fresh media was added to each well. Cells were cultured for one day prior to FACS to determine the percentage of cells with GFP [42]. The amount of virus used for the well with ≤30% transduced cells was selected to calculate the amount of virus required for the screen at MOI=0.3.

### 4.9 sgRNA library lentivirus production and transduction

A total of 1 × 10⁷ HEK293 cells were plated in each of the 150-mm dishes. The Lenti-eGFP/Lenti-gRNA:pVSVg:pGagPol (3:5:4) lentiviral packaging plasmids (36μg in total) were combined with 4.5mL Opti-MEM and 108 μL PEI for a period of 15 minutes. The cell culture medium was replaced, the mixture was added to the medium, and the cells were shaken gently and placed into the CO_2_ incubator. Following a period of 72 hours, the lentivirus-containing medium was collected and subjected to centrifugation at 5,000 rpm for a duration of three minutes at a temperature of 4°C. Thereafter, the medium was filtered through a 0.45 μM filter and dispensed and stored in a -80 refrigerator.

### 4.9 Generation of mutant cell libraries and screening

A total of 91,566 gRNAs were designed in the bovine whole genome library, and a total of 3×10^8^ MDBK-Cas9 cells were required according to the coverage of 1,000 and the multiplicity of infection (MOI) of infection=0.3. The 8×10⁶ allocation was distributed among 150-mm cell culture dishes in accordance with the aforementioned distribution plan. In order to prepare the lentiviral mixture solution, a volume of 50.4 mL of lentiviral solution should be mixed with 8 μg/mL polybrene. Subsequently, the mixture should be vortexed for 15 seconds, after which 0.92 mL of the resulting solution should be added to each plate. Following a 24-hour period of lentivirus infection, the medium should be changed to a standard D10 medium. Two days after the change of lentivirus, the medium should be changed to a drug medium (puro 2.2μg/mL), and the library cells should be digested and harvested after 10 days of cell drug screening.

### 4.10 Illumina sequencing of sgRNAs in the genome-wide library and enriched mutants

The surviving cells were employed for the extraction of the cellular genome through the utilisation of the Quick-DNA Miniprep Plus Kit (ZYMO RESEARCH #D4068). The gRNA region was amplified using Takara Ex Taq polymerase (Takara#RR001Q) and P7 and P5 primers carrying different barcodes. The amplified product was purified using the FastPure Gel DNA Extraction Mini Kit (Vazyme#DC301-01). The purified products were then submitted to Annoroad Gene Technology (Beijing) Co. for NGS sequencing. The sequencing results were analysed for gene enrichment using MaGeCk (Version 0.5.9) [20].

### 4.11 TCID50 assay

MDBK cells were seeded at a density of 2×10⁵ cells per well in 96-well plates and cultured for a period of 18 hours. The BPIV3 virus to be tested was thawed on ice and diluted at a 10-fold multiplicity of infection, with eight wells per dilution. The medium in the cell plates was aspirated, washed three times with PBS, and incubated with 100 μL of virus solution for one hour. The venom was aspirated, washed three times with PBS, and 200 μL of serum-free DMEM medium was added. The cell lesions were observed on the fourth day after infection. The cytopathic effects were recorded and median tissue culture infective dose (TCID50) were calculated using the Reed-Muench method as described previously [43].

### 4.12 Binding Entry assay

The non-target and indicator knockout cells were cultured at a density of 1x10⁵ in 6-well plates. Once the fusion of the cells had reached 80-90%, the cells were washed three times with phosphate-buffered saline (PBS) at 4°C. Subsequently, the cells were infected with BPIV3 at a multiplicity of infection (MOI) of 1 at 4°C for one hour. At the conclusion of the infection period, the plates were washed three times with PBS at 4°C. For the binding assay, the cells were subjected to direct digestion for subsequent qPCR analysis. For the entry assay, the cells were internalised at 37 °C for 30 minutes, after which the uninternalised virus was removed by washing with PBS at pH 3, and the cells were collected for qPCR. The aforementioned harvested cells were employed for the extraction of RNA using the Ultrapure RNA Kit (Cwbio#CW0581), and the RNA was subsequently reverse transcribed using the PrimeScript ‘FAST RT Reagent Kit with gDNA Eraser’ (Takara#RR092A). The RNA was reverse transcribed using the PrimeScript “FAST RT Reagent Kit with gDNA Eraser” (Takara#RR092A) and detected by qPCR on an ABI 7500 machine using TB Green® Premix Ex Taq™ II FAST GPCR (Takara#CN830A) qPCR dye and primers from the provided table S4.

### 4.13 Primary viral transcription and viral genome replication assays

To detect the initial transcription of the viral NP gene, the virus was initially infected in accordance with the aforementioned methodology, and RNA extraction was conducted three hours post-infection using the aforementioned method. For the purpose of reverse transcription, a PrimeScript™ RT reagent Kit with gDNA Eraser (Takara#RR047A) reverse transcription kit was utilised, employing specific reverse transcription primers in accordance with the manufacturer’s instructions (RT-vRNA:TGTTCGGAAATATGAATTTA, RT-cRNA:TCCTCTCCTATTTCTTCCCTGT, RT-mRNA:TTTTTTTTTTTTTTTTTTV). The aforementioned products were then subjected to qPCR experimentation, employing the aforementioned experimental procedures [44].

### 4.14 BPIV3 virus purification

The BPIV3 virus was concentrated using the Universal Virus Concentration Kit (Beyotime #C2901S), following the manufacturer’s instructions. To summarize, the viral supernatant was initially collected and cellular debris was removed through centrifugation and filtration. The pre-cooled Virus Precipitation Reagent was then mixed in proportion with the viral supernatant, thoroughly mixed, and stirred at a low speed at the appropriate temperature overnight. This was followed by centrifugation to remove the supernatant and careful aspiration of the precipitate. The process was then repeated, and the supernatant was collected as concentrated virus.

### 4.15 Confocal microscopy

For R18-labeled viruses, The concentrated and purified BPIV3 virus were co-labeled with 67 μmol/L R18 (Thermo Fisher #O246) in 1 mL total volume [29]. At the conclusion of the labelling process, any unbound R18 was filtered out using a 0.45 μm filter [29]. MDBK cells were subjected to three washes with PBS, after which the labelled virus was added and allowed to bind at 4°C for one hour. At the conclusion of the procedure, any unbound virus was removed by washing with PBS, and the cells were incubated at 37°C for the indicated times to allow for internalisation (early endosomes: 10 minutes, late endosomes: 20 minutes, lysosomes: 30 minutes). The cells were fixed using 4% paraformaldehyde for a period of 10 minutes. Subsequently, the cells were punched using Immunostaining Permeabilization Buffer with Saponin (Beyotime#P0095), in accordance with the instructions provided by the manufacturer. The second antibodies were used in a 1:500 configuration: anti-Rab5 (Immunoway#YT5456), Rab7 (Abcam#ab126712), and Lamp1 (Proteintech#65051). Cells were incubated overnight. Following the application of the anti-rabbit secondary antibody (Proteintech#SA00003) in a 1:200 configuration, the cells were incubated for a period of 2 hours. Subsequently, the cells were stained with Hoechst 33258 Staining Solution for 10 minutes. The images were captured using a Zeiss Lsm710.

### 4.16 Image Analysis and Colocalization

Images were analyzed using ZEISS ZEN 3.8 software. Colocalization analysis was performed on 30 regions of interest (ROIs) (6–10 ROIs per cell) across 3–5 cells per experimental group using the Coloc 2 plugin in Fiji. The frequency of association between Rab5, Rab7, and Lamp1 with R18-labeled virus was quantified using Manders’ Colocalization Coefficients. These coefficients range from 0 to 1, indicating the fraction of intensity in one channel that colocalizes with above-zero (or threshold) intensity in the other channel.

### 4.17 Lectin assays

Biotinylated Sambucus Nigra (SNA; Vector Biolabs#B-1305-2) and Maackia Amurensis Lectin II (MAL; Vector Biolabs#B-1265-1) were purchased from Vector Biolabs. Both control and primary bovine nasal turbinates, lung fibroblasts and SLC35A1 KO cells were fixed in 4% formaldehyde for 10 minutes. They were then washed twice with PBS. The cells were then incubated with 20 µg/mL biotinylated lectin for 1 hour at 4°C. The cells were then stained with 1 µg/mL Vari Fluor 594 streptavidin (MCE#HY-D1806). Lectin binding levels were then visualised using a confocal microscope.

### 4.18 Treatment with 3FAx-Neu5Ac

MDBK cells were treated with DMSO or 200µM 3FAx-Neu5Ac (MCE#HY-110288) for a period of 10 days to eliminate residual sialic acid from the cell surface. Lectin binding analysis was performed as described above.

### 4.19 Transmission electron microscope

MDBK non-target cells with LSM12 KO cells were seeded at a density of 5×10⁶ cells per 100 mm petri dish. BPIV3 was used to infect the cells at a multiplicity of infection (MOI) of 20 for a period of 40 min. At the conclusion of the infection period, the cells were fixed using 2.5% ultra-clear glutaraldehyde for a duration of 10 minutes. Thereafter, the cells were scraped using a cell scraper, subjected to centrifugation at 1000 rpm, and resuspended by replacing the cells with a new fixative. The transmission electron microscope images presented herewith were obtained from Servicebio.

### 4.20 Inhibitors assay

To investigate the effects of specific inhibitors on BPIV3 transport and their antiviral activities, the following experimental procedures were employed. Cells were pre-treated with the PIKfyve inhibitor apilimod (MCE#HY-14644), the NAADP blocker Ned-19 (MCE#HY-103316A), or the TRPML1 blocker (1R,2R)-ML-SI3 (MCE#HY-134819A) for 1 hour at 37°C.To assess the impact of these inhibitors on BPIV3 transport, MDBK cells were infected with BPIV3 and incubated at 4°C for 1 hour. The cells were subsequently washed with PBS and incubated with the respective inhibitor for an additional 30 minutes at 37°C. After washing with PBS, the cells were fixed with 4% paraformaldehyde for 10 minutes. Following incubation with the Lamp1 antibody to target the cells, confocal microscopy was used for observation. To investigate the antiviral effects of the inhibitors, cells were infected with BPIV3 at a multiplicity of infection (MOI) of 0.1 for 1 hour at 37°C. The respective inhibitor was added during the 24-hour incubation period. After incubation, the cell sediment and supernatant were collected from the plate for further analysis.

### 4.21 Cell viability assay

Cell viability was evaluated using the Cell Counting Kit-8 (CCK-8) assay. MDBK cells were seeded into 96-well plates at a density of 2000 cells per well in 100 µL of complete culture medium and incubated at 37°C overnight to ensure cell attachment and growth. Following pre-incubation, cells were treated with various concentrations of the respective inhibitors for the indicated time periods. Subsequently, 10 µL of CCK-8 solution was added to each well, and the plate was incubated for an additional 1.5 hours at 37°C. The absorbance was measured at 450 nm using a microplate reader (model number, manufacturer). Cell viability was calculated as a percentage of the control group using the following formula: Cell viability (%) = (OD of treated group / OD of control group) × 100.

### 4.22 Ned19 treatment and viral infection assay

Cells were seeded in 6-well plates at a density of 2 × 10⁵ cells/well. After 24 hours, cells were pre-incubated with 10 μM/mL Ned19 (Sigma-Aldrich) or an equivalent volume of DMSO (control) for 1 hour at 37°C. Subsequently, cells were infected with each virus at a multiplicity of infection (MOI) of 0.01 for 1 hour. Unadsorbed virus was removed, and cells were washed twice with PBS. Cells were incubated at 37°C for 24 hours. Supernatants were collected, and virus titers were determined by TCID_50_ (for ORFV and HSV) or plaque assays (for BoHV-1, H5N1/NV, and H5N1/COW) according to standard protocols. Each experiment was performed in triplicate.

### 4.23 Plaque assays

A549 or MDCK cells were infected with Influenza virus or HSV at MOI 0.5 in serum-free DMEM supplemented with 1% BSA and 1 μg/μL TPCK trypsin (Influenza virus add). 48 h post-infection, supernatant was collected and serial-diluted. Two hundred microliters of the diluted supernatant was used to infect MDCK cells on 6-well plates and the number of plaques were counted after 72 h. The virus titer was calculated in Plaque forming units (PFU)/mL.

### 4.24 Statistical analysis

The statistical analysis was conducted utilizing GraphPad Prism 9. The significance of the findings was determined through the use of a two-tailed unpaired Student’s t-test, a one-way or two-way analysis of variance (ANOVA), with a P-value threshold of <0.05, which was considered statistically significant and denoted with the following symbols: *P < 0.033, **P < 0.002, and ***P < 0.001. Values that were not deemed significant were designated as ns.

## Notes

### Competing Interest Statement

The authors have declared no competing interest.

